# Identification and structural basis of a *Chloroflexus* protein with homology to *Bacillus* quorum sensing-related prenyltransferase

**DOI:** 10.64898/2026.08.29.745113

**Authors:** Takashi Matsui, Sumika Inoue, Shunsuke Yanagimoto, Aira Kaneko, Reiichi Tago, Arisa Suto, Miho Odagi, Yoshio Kodera, Hiroyuki Morita, Ikuro Abe, Masahiro Okada

**Affiliations:** School of Science, Kitasato University, 1-15-1 Kitasato, Minami-ku, Sagamihara, Kanagawa, 252- 0373, Japan; Center for Disease Proteomics, School of Science, Kitasato University, 1-15-1 Kitasato, Minami-ku, Sagamihara, Kanagawa, 252-0373, Japan; Faculty of Chemistry and Biochemistry, Kanagawa University, 3-27-1 Rokkakubashi, Yokohama, Kanagawa, 221-8686, Japan; Institute of Natural Medicine, University of Toyama, 2630-Sugitani, Toyama, Toyama, 930-0194, Japan; Faculty of Pharmaceutical Sciences, The University of Tokyo, 7-3-1 Hongo, Bunkyo-ku, Tokyo, 113-0033, Japan

## Abstract

Quorum sensing in Gram-positive bacteria commonly relies on posttranslationally modified peptide pheromones. In *Bacillus subtilis*, the prenyltransferase ComQ catalyzes tryptophan prenylation of the quorum-sensing peptide ComX, but the structural basis of this unique peptide modification has remained unclear. Here we identified a previously uncharacterized ComQ homolog, StheQ, and its cognate peptide substrate, StheX, from *Sphaerobacter thermophilus* and investigated their structural and functional relationship. Liquid chromatography-tandem mass spectrometry (LC-MS/MS) analysis demonstrated that StheQ catalyzes prenylation of the tryptophan residue located second from the C-terminus of StheX. Crystal structures of apo StheQ and its complexes with a farnesyl pyrophosphate analog revealed that StheQ adopts the all-α-helical fold of the trans-isoprenyl diphosphate synthase (IPPS) superfamily while possessing an active-site architecture adapted for peptide-based indole prenylation. The structures identified a single Mg^2+^-binding site associated with the first aspartic acid-rich motif and showed no evidence for metal coordination at the pseudo-second aspartic acid-rich motif. Site-directed mutagenesis, complex formation assays, and docking analyses identified a peptide-binding pocket adjacent to the active site and suggested that N215 contributes to productive positioning of the acceptor tryptophan. These findings establish the structural basis for peptide prenylation by a ComQ-family enzyme, providing insight into the evolution of peptide-based indole prenylation within the IPPS superfamily, and support the view that ComQ-family enzymes constitute a distinct functional branch specialized for peptide modification.

## Introduction

Quorum sensing is a bacterial cell-cell communication process mediated by extracellular signaling molecules and is a widespread phenomenon in bacteria.^1–3^ As the molecules accumulate in the environment, quorum sensing system stimulates the cell density-driven genetic regulations, such as bioluminescence, the formation of biofilms, antibiotic production, and competent cell formation. Gram-positive bacteria generally use oligopeptides, which are called quorum-sensing pheromones, as the extracellular signaling molecules encoded as precursors and are diverse in sequence and structure.^4^ In their biosynthesis, the precursor peptides are processed, and the mature pheromones frequently undergo posttranslational modifications.^4–6^

Prenylation is a broadly distributed post-translational modification in peptide natural products^7–10^ and has also been identified in quorum-sensing pheromones. One representative case is the tryptophan prenylation followed by intramolecular cyclization of the quorum-sensing pheromone ComX.^11,12^ ComX pheromones, encoded by the *comQXPA* gene cluster (Figure S1A), are found in *Bacillus subtilis* and related *Bacilli* and exhibit sequence diversity, ranging from 6 to 10 amino acids in length without a defined consensus sequence.^13,14^ Within this gene cluster, ComQ catalyzes the geranylation or farnesylation of the tryptophan residue of ComX in the presence of geranyl or farnesyl diophosphate (GPP or FPP) and Mg^2+^ ions^9,15,16^, thereby functioning as metal-dependent tryptophan prenyltransferase. While ComQ-mediated post-translational prenylation was initially thought to be unique to *Bacilli*, comprehensive analyses of gene arrangements have revealed that *comQXPA* loci are widely distributed across the phylum *Firmicutes*, including the class *Clostridia*.^17^ These findings suggest that ComQ-mediated prenylation may occur in more diverse taxa than previously recognized.

Although ComQ belongs to the trans-isoprenyl pyrophosphate synthases (IPPS) superfamily, prenylation of the indole C3 position at the tryptophan residue instead of isopentenyl diphosphate (IPP) condensation reaction, represents a chemically distinct reaction compared with common isoprenoid biosynthesis.^18^ Notably, prenyltransferases in this family exhibit broad substrate specificity and do not require a defined recognition sequence, allowing the modification of diverse tryptophan-containing peptides and their derivatives.^19,20^ Our recent research revealed the structure and catalytic mechanism of the all-α-helical indole prenyltransferase Ord1, which also belongs to the IPPS superfamily, providing a structural framework for aromatic prenylation within the IPPS superfamily.^21^ However, because no structure of a ComQ-family enzyme has yet been reported, it remains unclear whether these enzymes employ a similar catalytic strategy, leaving the structural basis of peptide-based indole prenylation unresolved.

In this study, we found a putative homolog of ComQ, which is called StheQ from *Sphaerobacter thermophilus* DSM 20745. *S. thermophilus* belongs to the phylum *Chloroflexi*, which is evolutionarily distinct from the phylum *Firmicutes*. Although StheQ and ComQ share only 19% sequence homology, the amino acid residues essential for ComQ’s enzymatic activity, including those forming the first aspartic acid-rich motif (FARM; DDXXD, X represents the amino acids that vary among the different strains) and the pseudo-second aspartic acid-rich motif (pseudo-SARM; NDXXX), are well conserved. (Figure S1B) In addition, immediately adjacent to the *stheQ* gene, there is an ORF encoding a 69-amino acid residue tryptophan-containing protein. Although this ORF exhibits low homology to ComX (Figure S1C), it shares a common feature: a short-chain peptide containing a C-terminal tryptophan residue, thus we designated it *stheX* and its encoded protein StheX.

Here, we report the crystal structures of StheQ in apo and ligand-bound forms and demonstrate StheQ functions as a peptide prenyltransferase similar to *Bacillus* ComQ. Combined with mutational and docking analyses, our results provide structural insight into how ComQ-family enzymes accommodate tryptophan-containing peptide substrates and coordinate prenyl donors, as well as into the evolution of peptide-based indole prenylation within the IPPS superfamily.

## EXPERIMENTAL PROCEDURES

### Expression and purification of StheQ variants for the activity assay

Codon-optimized gene encoding StheQ was synthesized and purchased from Eurofins Genomics (Ebersberg, Germany). The DNA fragment was cloned into a pET22-based vector to prepare a plasmid encoding TEE(MNHKV)-His6-Thrombin(SLVPRGS)-StheQ fusion protein (THT-StheQ). Plasmids for StheQ mutants were constructed by PCR-based site-directed mutagenesis using an In-Fusion HD Cloning Kit (TaKaRa Clontech, Shiga, Japan) based on the manufacturer’s procedure. The plasmid encoding THT-StheQ was used as a template for PCR. Primers for the PCR are listed in Table S1, and the sequences of the plasmids were confirmed by Sanger sequencing.

StheQ variants were overexpressed in *E. coli* DynaCompetent Cells Zip BL21 (DE3). (BioDynamics Laboratory Inc., Tokyo, Japan). Cells were grown at 37°C in LB medium containing 100 mg/L ampicillin and 0.5% (w/v) glucose until the optical density reached 0.6. 1 mM isopropyl β-D-1-thiogalactopyranoside (IPTG) were then added, and the growth was continued for 18 h at 20°C. The cells were harvested and resuspended in lysis buffer (31 mM Tris-HCl, pH 8.0, and 80 µM MgSO4) containing 0.1% *n*-dodecyl-β-maltoside. The cell lysate was sonicated and then centrifuged at 12,000×*g* for 30 min at 4°C. The supernatant was filtered and loaded onto a HisTrap HP column (Cytiva, Marlborough, USA) equilibrated with lysis buffer. The column was then washed with lysis buffer, and the variant proteins were eluted from the column by elution buffer (20 mM Na3PO4-HCl, pH 7.6, 150 mM NaCl, and 0-250 mM imidazole). The StheQ was concentrated using Amicon Ultra 10k centrifugal filter (Merck, Darmstadt, Germany) and suspended in buffer containing 50 mM HEPES-NaOH, pH 8.0, and 5 µM MgCl2. Protein concentrations were determined using NanoVue Plus (Biochrom, Cambridge, UK) based on Warburg-Christian method. The purified proteins were flash-frozen and stored at –80°C.

### Expression and purification of StheQ variants for crystallization and SEC analysis

StheQ was overexpressed in *E. coli* Rosetta2 (DE3) (Merck) harboring the pET-26b plasmid (Marck) containing the full-length *stheQ* cDNA. Cells were grown at 37°C in LB medium containing 100 mg/L ampicillin and 34 mg/L chloramphenicol until the optical density reached 0.6. 1 mM IPTG was then added to express C-terminal His6-fused StheQ and growth was continued for 16 h at 25°C. SeMet-labelled StheQ was overexpressed in *E. coli* Rosetta2 (DE3). Cells were grown at 37°C in M9 medium containing 1 mM MgSO4, 0.1 mM CaCl2, 1× BME vitamins solution (Merck), 0.4%(w/v) glucose, 100 mg/L ampicillin, and 34 mg/L chloramphenicol until the optical density reached 0.6. 25 mg/L selenomethionine (FUJIFILM Wako Pure Chemical Co., Osaka, Japan), 100 mg/L each of L-Lys, L-Thr, L-Ile, L-Leu, L-Val, and L-Phe, and 1 mM IPTG were then added to express SeMet-labeled and C-terminal His6-fused StheQ and growth was continued for 16 h at 25°C. All subsequent procedures were performed at 4°C. The cells were harvested and resuspended in lysis buffer (50 mM Tris-HCl, pH 8.0, 200 mM NaCl, 5 mM MgCl2, 5% glycerol, 1 mg/ml lysozyme, and 5 U/ml Benzonase nuclease (Merck)). The cell lysate was sonicated and then centrifuged at 10,000×*g* for 20 min at 4°C. The supernatant was loaded onto a Ni Sepharose 6 Fast Flow column (Cytiva). StheQ eluted from the column by elution buffer (50 mM HEPES-NaOH, pH 7.0, 100 mM NaCl, 5% glycerol, and 600 mM imidazole). The eluate was diluted fivefold with buffer A (50 mM HEPES-NaOH, pH 7.0, 5 mM MgCl2, and 2 mM DTT) and purified on the Resource Q column (Cytiva) using a linear gradient of buffer A to buffer B (50 mM HEPES-NaOH, pH 7.0, 1 M NaCl, 5 mM MgCl2, and 2 mM DTT). Finally, StheQ was concentrated to 2 mg/ml using Macrocep 10k centrifuge device (PALL, Port Washington, USA).

All mutants of StheQ were constructed with a Quickchange site-directed mutagenesis kit (Stratagene, La Jolla, USA), according to the manufacturer’s protocol. Primers for the PCR are listed in Table S2, and the sequences of the plasmids were confirmed by Sanger sequencing. The mutants were expressed and purified by the same procedure as used for the wild-type enzyme for crystallization and SEC analysis.

### Protein purification of StheX

StheX was overexpressed in *E. coli* Rosetta2 (DE3) (Merck) harboring the plasmid modified pET-22b (Merck) containing full-length *stheX* cDNA. Cells were grown at 37°C in LB medium containing 100 mg/L ampicillin and 34 mg/L chloramphenicol until the optical density reached 0.7. 1 mM IPTG was then added to express N-terminal His6 and TEV protease recognition sequence-fused StheX and growth was continued for 4 h at 30°C. The purification of StheX was performed using a previous procedure.^22^ The cells were harvested and resuspended in denatured buffer (50 mM Tris-HCl, pH 8.0, 200 mM NaCl, 5% glycerol, and 6 M guanidine-HCl). The cell lysate was sonicated and then centrifuged at 10,000×*g* for 20 min at 4°C. The supernatant was loaded onto a Ni Sepharose 6 Fast Flow column (Cytiva). Refolding of the precursor was performed using a linear gradient of denatured buffer to refolded buffer (denatured buffer without guanidine-HCl) and eluted from the column by elution buffer (50 mM HEPES-NaOH, pH 7.0, 100 mM NaCl, 1 mM DTT, and 600 mM imidazole). The elution was purified on the HiLoad 16/60 Superdex75 (Cytiva) using storage buffer (50 mM HEPES-NaOH, pH 7.0, 100 mM NaCl, 5 mM MgCl2, and 2 mM DTT). Finally, the StheX was concentrated to 1.7 mg/ml using Macrocep 3k centrifuge device (PALL).

### Identification of the prenylation in StheX

To identify the prenylation of StheX, 40 µM StheQ was mixed with 40 µM StheX and 1 mM FPP in 25 µL of 100 mM HEPES-NaOH pH 7.0 and 5 mM MgSO4 and incubated at 37°C for 16 h. The reaction mixture was incubated with 24 mM Bond-Breaker TCEP solution (Thermo Fisher Scientific) at 80°C for 10 min and then alkylated by 33 mM iodoacetamide at room temperature for 30 min in the dark for the protection at the thiol group of Cys residue. Alkylated protein samples were mixed with 20 µL of Sera-Mag SpeedBeads (hydrophilic:hydrophobic particles= 1:1, Cytiva) suspension to remove alkylating reagents followed by previous study.^23^ The beads were resuspended in 100 µL of Tris-HCl pH 8.0 containing 200 ng Lys-C and 200 ng Trypsin and then the sample was digested at 37°C for overnight. The magnetic beads were removed using a magnetic stand, and the resulting digested peptides were loaded onto a MonoSpin C18 column (GL Science, Tokyo, Japan) for desalting. The digested peptides were eluted with 100% CH3CN containing 0.1% formic acid. The elution was diluted to 50% CH3CN containing 0.1% formic acid. the diluted sample was injected into an online LC-MS/MS system with a Vanquish Neo UHPLC system (Thermo Fisher Scientific) connected to an Orbitrap Exploris 240 mass spectrometer (Thermo Fisher Scientific). The digested peptides were eluted with a gradient of water and 80% CH3CN, both containing 0.1% formic acid: 0–10 min, 36%–76% CH3CN, on an analytical column (C18, particle diameter 3 µm, 75 µm × 125 mm; Nikkyo Technos, Tokyo, Japan) at a flow rate of 300 nL/min. The analytical conditions were as described previously^24^ with slight modifications. MS spectra were collected over the scan range of 750–1000 *m*/*z* at 90,000 resolution and a maximum injection time of auto. The 10 most-intense ions with charge state of 2^+^ that exceeded an intensity of 2 ×10^4^ were fragmented by collision-induced dissociation with normalized collision energies of 20, 25, and 30%. MS/MS spectra were acquired on the Orbitrap mass analyzer with a mass resolution of 60,000 to reach an AGC target of 1 ×10^5^.

LC-MS/MS data were analyzed using PEAKS Studio 13 build 20250715 (Bioinformatics Solutions Inc., Waterloo, Canada) Database searching was implemented against the UniProt protein sequence database (*E. coli*, proteome ID: UP000000625) supplemented with StheQ and StheX sequences. The following parameters were applied: enzyme, semi-specific trypsin; precursor mass tolerance, 10 ppm; fragment ion tolerance, 0.02 Da; Variable modification, Trp farnesylation with cyclization (+204.19), Cys carbamidomethylation (+57.02), and Met oxidation (+15.99). Peptide identification was filtered at a false discovery rate <1%.

### StheQ enzyme assay using Fmoc-Trp and StheX(60-69) as substrates

To assess prenylation activities of StheQ and its mutants, 10 µL of the solutions containing 2 mM FPP, 5 mM MgCl2, 50 mM HEPES-KOH (pH 8.0), 25 µM StheQ variants, and 100 µM substrates (Fmoc-Trp or StheX(60-69), a synthetic peptide corresponding to the C-terminal 10 residues of StheX) were prepared and incubated at 37°C for 3 h. After the incubation, 10 µL water and 20 µL methanol were added, and the solutions were incubated on ice for 5 min. Then, the solutions were centrifuged (11,000×*g* for 1 min), filtered, and the supernatants were analyzed by liquid chromatography-mass spectrometry (LC-MS). The StheX(60-69) was synthesized and purchased from Eurofins Genomics. The authentic standards for LC-MS analysis were chemically synthesized according to a previously reported procedure.^25^

LC-MS measurements were conducted using a Xevo G3-XS QToF instrument connected to an Acquity H-Class UPLC system (Waters, Milford, USA). All analyses were conducted in positive mode with a capillary voltage of 1.5 kV. Nitrogen was used as both the cone gas (flow rate: 60 L/min) and desolvation gas (flow rate: 1000 L/min). The ionization source was maintained at 150°C, while the desolvation gas was set to 500°C. Samples containing Fmoc-Trp were analyzed using a InertSustain C30 column (100Å, 3 µm, 2.1 mm × 50 mm, GL sciences, Tokyo, Japan) maintained at 30°C with a linear gradient of 70–98% solvent B, where solvent A consisted of 0.05% formic acid in water and solvent B consisted of 0.05% formic acid in methanol. Samples containing StheX(60-69) were analyzed using a InertSustain C18 column (200Å, 2 µm, 2.1 mm × 50 mm, GL sciences) maintained at 40°C with a linear gradient of 10–98% solvent B, where solvent A consisted of 0.05% formic acid in water and solvent B consisted of 0.05% formic acid in acetonitrile. Data analysis was carried out using MassLynx 4.2 software (Waters), and to generate extracted ion chromatograms (EICs) for specific ions, the *m*/*z* tolerance window was fixed to 0.02. The conversion efficiencies were calculated based on the EIC peak area of unmodified substrates and farnesylated products. MS/MS spectrum was acquired with a 0.1 s scan time, with 20-40 eV collision energy using MSe mode.

### Crystallization and Data collection

Well-diffracted StheQ apo crystal was obtained using 100 mM MES-NaOH, pH 6.8 containing 10%(w/v) PEG 8000 and 10 mM L-Ile with sitting-drop vapor-diffusion method by mixing 0.5 µL of the purified StheQ with an equal volume of reservoir solution and equilibrating the mixture against 50 µL reservoir solution after a few days at 5°C. To obtain binary complex crystal of StheQ with FSPP, StheQ was crystallized using 100 mM MES-NaOH, pH 6.8 containing 10%(w/v) PEG8000 and 10 mM glycyl-glycyl-glycine with sitting-drop vapor-diffusion method by mixing 0.5 µL of the purified StheQ with an equal volume of reservoir solution and equilibrating the mixture against 50 µL reservoir solution after a few days at 5°C. To determine the metal binding site, StheQ was crystallized using 100 mM MES-NaOH, pH 6.6 containing 10%(w/v) PEG8000 and 10 mM L-Asn with sitting-drop vapor-diffusion method by mixing 0.5 µL of the purified StheQ with an equal volume of reservoir solution and equilibrating the mixture against 50 µL reservoir solution after a few days at 5°C. For structure determinations of StheQ with Mg^2+^ and farnesyl thiodiphosphate (FSPP, an FPP analog, Echelon Biosciences, Inc. Salt Lake City, USA), and of metal binding site were soaked into the reservoir solutions including 10 mM FSPP and 10 mM MgCl2 for 5 h and 5 mM YbCl3 for a few min, respectively. These crystals were transferred into a cryoprotectant solution, containing of each crystallization solution with 20%(v/v) glycerol. StheQ A167Y mutant crystal was obtained using 100 mM sodium acetate pH 4.5 containing 15%(v/v) isopropanol with sitting-drop vapor-diffusion method by mixing 0.5 µL of the purified A167Y mutant with an equal volume of reservoir solution and equilibrating the mixture against 50 µL reservoir solution after a few days at 5°C. The A167Y mutant crystal was transferred into a cryoprotectant solution, containing of the crystallization solution with 25%(v/v) ethylene glycol. After a few seconds, the crystal was picked up in a nylon loop and then flash-cooled at −173°C in a nitrogen-gas stream.

Single-wavelength anomalous diffraction (SAD) data of SeMet-labeled StheQ apo and the crystal soaked into YbCl3 solution, and diffraction data of complex crystal of StheQ-FSPP and A167Y mutant crystal were collected on beamline PF-AR NW-12A, BL-17A, BL-5A, and NW12A at Photon Factory, Tsukuba, Japan under cryogenic conditions at 100K, respectively. For structure determinations of SeMet-labeled StheQ and StheQ-Yb complex, Wavelengths of 0.97934 and 1.24198 Å were used for data collection based on the fluorescence spectrum of the selenium (Se) *K* and ytterbium (Yb) *L* absorption edges, respectively.^26^ The diffraction data was processed and scaled using *XDS*. Se and Yb sites were determined and refined, and initial phase of StheQ apo was calculated with *AutoSol* in *PHENIX.*^27^ Molecular replacements (MRs) of StheQ-FSPP and A167Y mutant were performed with StheQ as the search model using *Molrep* in the CCP4 suite.^28^ The structures were rebuilt using *AutoBuild* in *PHENIX*,^27^ modified manually with *Coot*^29^ and refined with *PHENIX.*^27^ StheQ-Yb complex and Yb sites were determined by MR-SAD method calculated with *AutoSol* in *PHENIX*. All crystallographic figures were prepared using ChimeraX.^30^

### SEC analysis of enzyme–peptide complex formation

The reaction mixture contained 3.0 nmol of StheQ and 3.0 nmol of His-StheX in reaction buffer (50 mM sodium phosphate buffer, pH 6.4 and 5 mM MgCl2). Incubations were performed at 20°C for 30 min and mixture was loaded onto the Superdex200 10/300 size-exclusion column (Cytiva) equilibrated with reaction buffer. The peaks were collected and were analyzed by Tricine SDS-PAGE.^31^

### Modeling and docking studies

Three-dimensional models of the StheX(60-69) was generated and optimized using CNS 1.2 ^32^. The StheQ model was built by swapping the α7 helix in molecule A of StheQ-FSPP with α7 helix in molecule B of StheQ-FSPP. The StheX(60-69) was then docked into the above model by Autodock Vina 1.0.2 ^33^. The three-dimensional structure of StheX (1-69 residues) was also predicted using ColabFold^34^.

## RESULTS

### Enzymatic property of StheQ

A shorter ORF was located immediately downstream of *stheQ* in the same gene cluster and was predicted to encode a 69^-^residue protein containing two tryptophan residues at the second and third positions from the C-terminus (Figure S1B). This ORF was designated *stheX* and its encoded protein was named as StheX. To determine whether the putative cognate peptide StheX serves as a substrate of StheQ, full length of StheX was incubated with StheQ, and the reaction mixtures were analyzed by LC-MS/MS. MS/MS spectra of StheX revealed that the tryptophan residue at the second position from C-terminus was modified by prenylation (Figure S2A). These findings identify StheX as the cognate peptide substrate of StheQ and establish StheQ as a ComQ-like peptide prenyltransferase.

To further validate modification of the tryptophan residue, enzymatic activity of StheQ was evaluated using Fmoc-Trp as a model substrate. UPLC analysis revealed the formation of a reaction product with the same retention time as the chemically synthesized C3-α-farnesylated Fmoc-Trp (Figure 1). Furthermore, the MS/MS fragmentation pattern of the enzymatic product was consistent with that of the authentic standard, further supporting StheQ-mediated farnesylation of tryptophan residue (Figure S2B).

**Figure 1.**
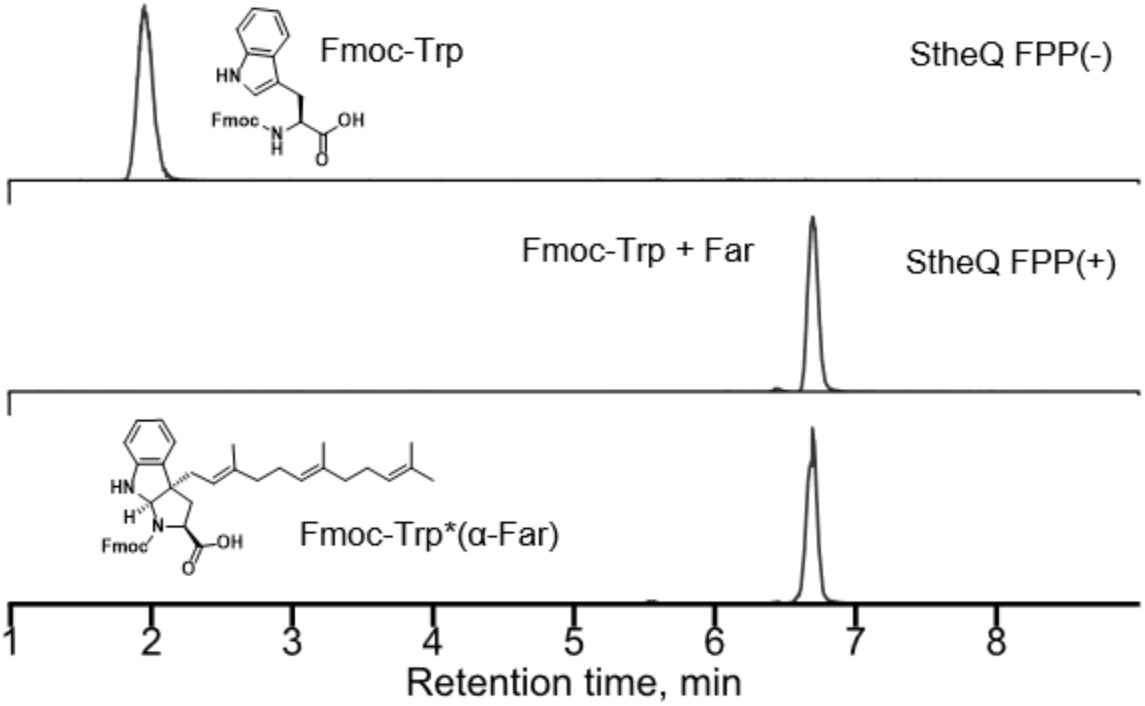
Substrate acceptance of StheQ toward Fmoc-Trp with and without FPP (Far: farnesyl). The authentic standard of α-farnesylated Fmoc-Trp is included for comparison. Combined EICs of *m*/*z* 427.165 and 631.353 are shown.

### Overall structures of StheQ

Based on the enzymatic assays, StheQ exhibits ComQ-like tryptophan prenyltransferase activity. To further elucidate its catalytic mechanism, the crystal structure of StheQ apo was determined at 2.43 Å resolution using Se SAD method (Figure 2A). The crystals contained a homodimer in the asymmetric unit. A slight structural difference was observed between molecules A and B with root mean square deviations (rmsd) value of 0.43 Å (Figure 2B). The residues 228-259 in each molecule were disordered except the residues 230-241 in molecule A. In the molecule B, the residues of α6-α7 loop (residues 156-164), residues 213-227, and 260-266 of StheQ apo were also disordered (Figure 2B). In the molecule A, a deep and wider cavity was located on the center of the protein. This cavity was formed by α2-α3 loop, α3, α4, α6, α6-α7 loop, α7 α8, α9, and α10 (Figure 2C). The residues derived from C-terminus His6-tag and restriction enzyme sequences were also ordered their structures and interacted with α8 and α9 helices.

**Figure 2.**
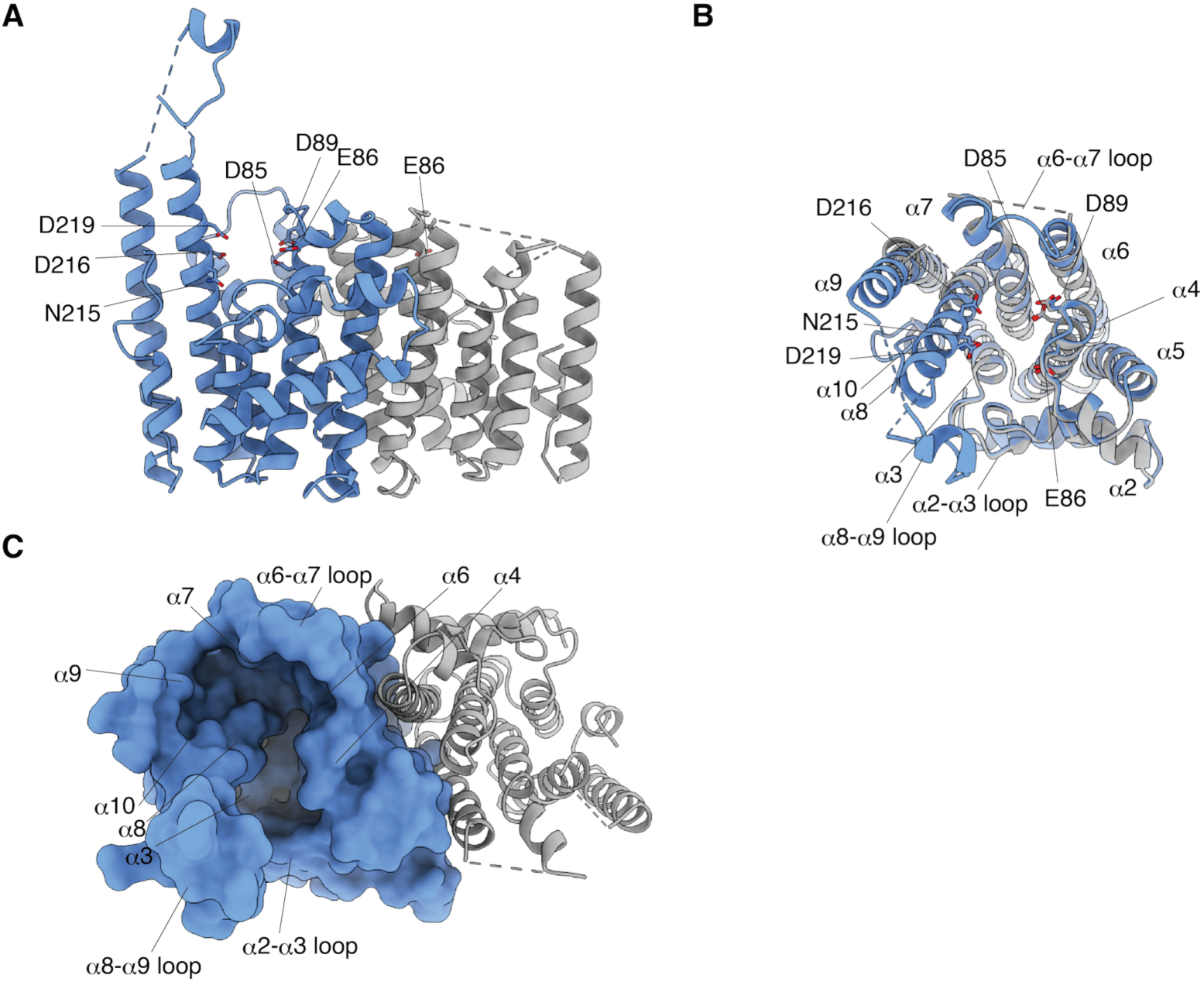
Overall structure of StheQ. (A) Dimer conformation in asymmetric unit. (B) Each subunit was superimposed. Molecules A and B are shown in slate and gray colors, respectively. Side chains of residues in the FARM and pseudo-SARM are shown as stick models. (C) Surface representation of StheQ molecule A. Deep pocket is composed of α2– α3 and α6– α7 loops, and α3, α4, α6, α7, α8, α9, and α10 helices.

A structure-based similarity search using the *Dali* program^35^ revealed that the overall structure of StheQ exhibited rmsd values of 2.6 and 3.5 Å to geranylgeranyl pyrophosphate synthase (GGPPS, PDB entry 3PDE, z-score 25.1, 19% sequence identity)^36^ and indole prenyltransferase (Ord1, PDB entry 9V4J, z-score 25.2, 19% sequence identity)^21^, respectively (Figures 3 and S3). Therefore, the structure of StheQ confirmed the all α-helical prenyltransferase fold as seen in the structures of IPPS and all-α-helical indole transferase, and StheQ belongs to the IPPS superfamily (Figure 3). Notably, although StheQ exhibits comparable overall structural similarity to both GGPPS and Ord1, the relevance of Ord1 as a functional reference becomes evident when considering substrate chemistry. Unlike canonical IPPS enzymes such as GGPPS, which act as the elongation of isoprene unit to generate longer isoprenyl pyrophosphate, Ord1 catalyzes aromatic indole prenylation. Given that StheQ modifies the indole side chain of a tryptophan residue within a peptide substrate, structural similarity at the level of the prenyl acceptor-binding pocket and surrounding catalytic residues is more informative than global fold similarity. Although Ord1 also has a lid region and formed closed conformation accompanied with the occupation of the pocket, opening the lid region in StheQ may guarantee to access to the pocket formed by α6-7 loop, α7, α9 and α10 helices, and thus this pocket is likely to be important for the production of the prenylated peptide and/or the interaction of StheX (Figure S3C–F).

**Figure 3.**
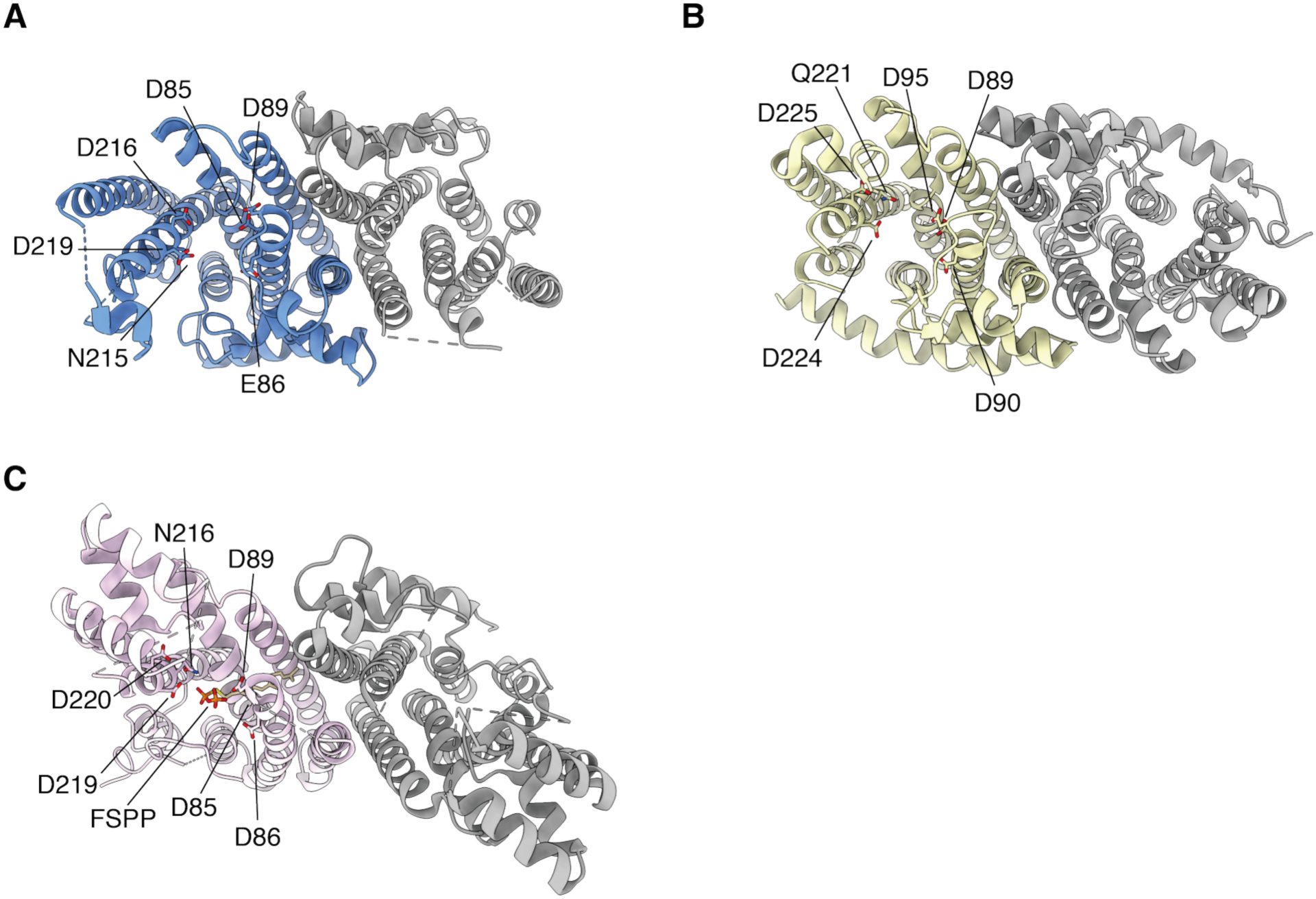
Structure comparisons of StheQ apo with other proteins belonging to all-α helical prenyl diphosphate synthase superfamily. Crystal structures of (A) StheQ, (B) GGPPS (PDB 3PDE), and (C) Ord1 bound with FSPP (PDB 9V4J). Molecule A in StheQ apo, GGPPS, and Ord1 are indicated as slate, yellow and pink cartoons, respectively. Molecule B of each enzyme is shown as gray cartoon. The side chains of residues in FARM, SARM, and pseudo-SARM are shown as stick models.

### Architecture of FPP binding site

IPPS enzymes have two conserved aspartic acid-rich motifs: FARM and SARM. The aspartic acid-rich motif composed of the conserved amino acid sequence DDXXD. StheQ not only shares the residues in the conserved FARM sequence but also has one pseudo-SARM instead of SARM. FARM (D85, E86, and D89) and pseudo-SARM (N215, D216, and D219) in molecule A of StheQ apo located in α4 and α8 helices, respectively, and faced each other at both side of the wall of the cavity (Figure 2). However, the conformation of the FARM was ordered in each subunit, in contrast, the part of the α8 helix including pseudo-SARM in molecule B was disordered in the crystal structure.

To understand the recognition mechanism of StheQ with FPP, the complex structure of StheQ with FSPP, which is a mimic of the FPP, was solved at 2.47 Å resolution (Figure 4A). Asymmetric unit contained a homodimer as seen in the crystal of StheQ apo. Structural comparison between apo- and FSPP bound-forms exhibited that the conformations of molecule A and B in StheQ apo are very similar to those in StheQ-FSPP (rmsd values of 0.30–0.45 Å) (Figure 4B). However, residues 226-260 in each subunit of StheQ-FSPP were disordered as seen in the structure of molecule B in StheQ apo. The farnesyl moiety was located in a similar manner in each subunit and was accommodated into the prenyl-moiety binding tunnel (Figures 4C–D and S4). An alternative conformation of the pyrophosphate moiety was also observed (Figure S4) and occupied a position composed of α2–α3 loop, α4, and α8 helices as the seen in the acceptor site of polyprenyl synthase from *Caulobacter crescentus* (PDB 3OYR).^36^ The pyrophosphate in the molecule A and the other alternative position of molecule B was located the similar position composed of FARM (D85, D89, and D91), K174, and Q212 at one helical pitch forward to pseudo-SARM (Figures 4C and S4), and was shared position as seen in the case of FSPP complex structure of an indole prenyltransferase Ord1 (Figure 4D). In particular, Q212 formed hydrogen bonds with α phosphate as seen in N215 of Ord1 which is a key residue involved in the prenylation of the prenyl acceptor substrate.^21^ IPPS enzymes, such as FPPS (PDB 1RQI), exhibited that two Mg^2+^ ions and one Mg^2+^ ion are coordinated with FARM and SARM, respectively.^37^ In the Ord1 structure, two Mg^2+^ ions also coordinated with FARM residues,^21^ and thus the conserved DDXXD motif was involved in the recognition of prenyl donor molecule and the catalysis of isoprenoid transfer via Mg^2+^ ion.^21,37,38^

**Figure 4.**
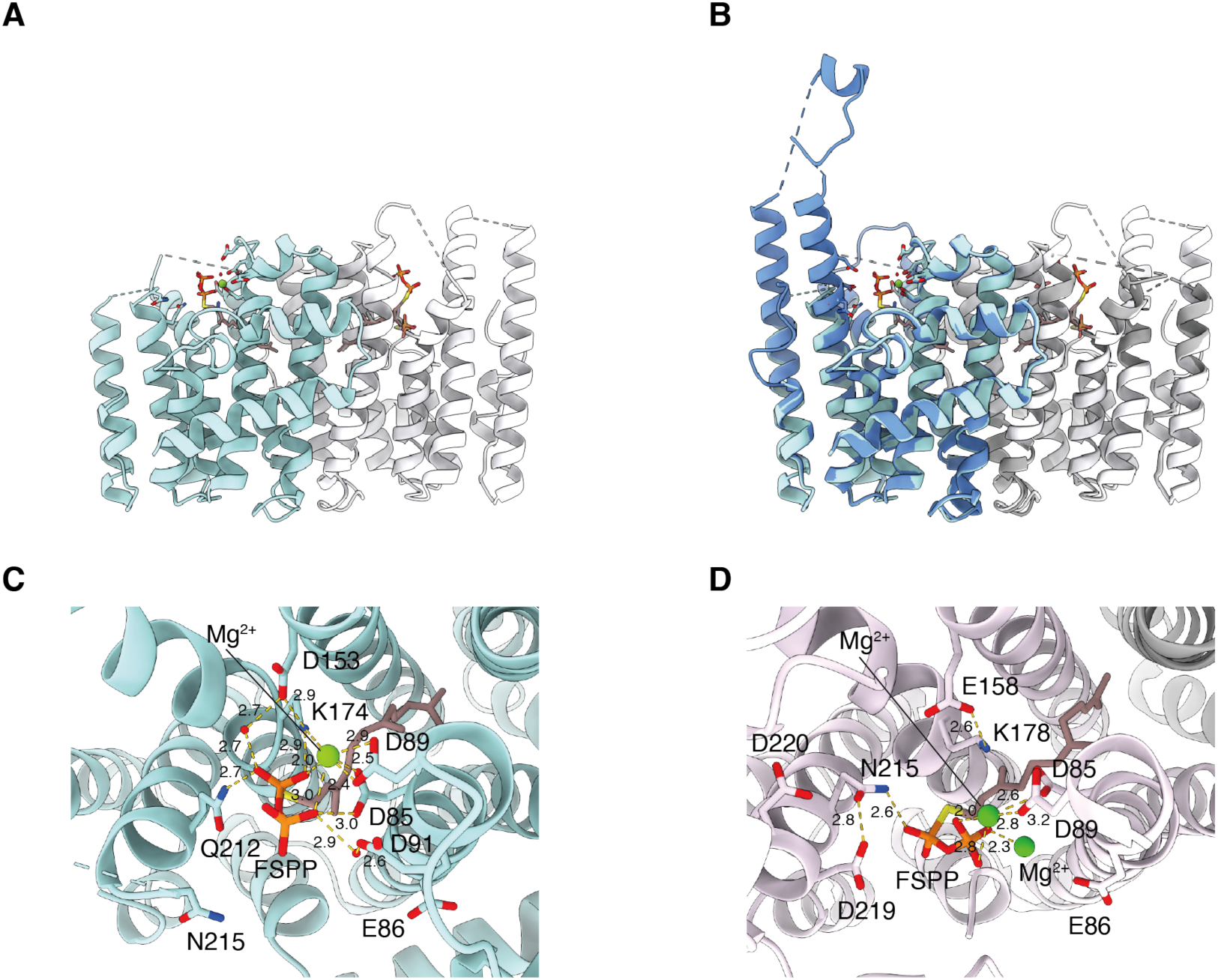
Complex structure of StheQ bound with FSPP. (A) Overall structure of StheQ bound with FSPP. The two protomers are shown as cyan and white cartoons, respectively. (B) Superimposing of StheQ–apo with StheQ-FSPP form. (C, D) Closed-up view of the region around FARM and pseudo-SARM in (C) StheQ and (D) Ord1. Residues in FARM and pseudo-SARM and those interacting with FSPP are shown as stick models. Mg^2+^ coordinations and hydrogen bonds are shown as yellow dotted lines.

In the StheQ-FSPP complex, the metal ion was located at the FARM and coordinated with the pyrophosphate, D85, and D89, was assigned as Mg^2+^ ion. To probe potential metal-binding sites, we determined the structure of StheQ soaked with YbCl3 at 2.71 Å resolution using Yb-SAD phasing, with Yb^3+^ serving as an anomalous scatterer. The Yb^3+^ site overlapped with the metal site observed in the StheQ-FSPP complex, supporting the assignment of the FARM-associated site as a Mg^2+^-binding site^25^ (rmsd values of 0.3 Å, Figure S5A–C). The *F*o – *F*c map at 5.0 σ showed a clear Yb peak in each subunit at coordination distances to D85 and D89 in FARM within 2.5 Å, respectively (Figure S5D–G).^39^ One of the Yb^3+^ ion in StheQ-Yb complex was placed on the similar position as seen in the metal ion in StheQ-FSPP (Figure S5B). Although the coordination of two Mg^2+^ ions with residue in the FARM were observed among with StheQ, Ord1^21^, and IPPS^37^, no Yb^3+^ ion appeared in the *F*o – *F*c map at 5.0 σ around the pseudo-SARM region despite the coordination of Mg^2+^ ion with SARM residues in other IPPS enzymes. These results predicted that only the residues in FARM via Mg^2+^ ion coordination and Q212 at one helical pitch forward to pseudo-SARM via hydrogen bond recognized and catalyzed the FPP molecule as the same manner observed in Ord1.^21^

### Interaction mechanism of StheQ with StheX

Previous investigation reported that N186D mutant enzyme of ComQRO-E-2, which corresponding with N215 in StheQ, exhibited approximately a 90% reduction of ComX-forming activity *in vitro*.^40^ The previous investigation may indicate that residues in the pseudo-SARM participates in either or both the transfer of the isoprenoid and/or interaction with StheX. To identify the peptide recognition mechanism, we also constructed site-directed mutants in pseudo-SARM to assess the effect of the interaction between mutant and StheX, since StheQ strongly interacts with StheX *in vitro* and complex of StheQ with StheX was detected by the size-exclusion chromatography (SEC) using a Superdex 200 10/300 column (Figure 5A). The complex peak was eluted as same retention volume as wild type using N215D, D216A or D219A in pseudo-SARM as a binding partner. These results revealed that N215, D216, and D219 are not crucial for the binding of StheX. Therefore, to further clarify the peptide binding site, we also constructed A167Y and T267L mutants located at α7 and α9 helices, respectively. Although the complex peak of T267L mutant with StheX was slightly delayed suggesting the weaker interaction of T267L mutant with StheX than that of wild type, no complex peak appeared using A167Y mutant and the individual peaks of A167Y mutant and StheX were observed (Figure 5A). These SEC analyses together with the StheQ structure, these observations identified a candidate pocket comprising α6-α7 loop, and α6, α7, α8, and α9 helices for StheX recognition. To evaluate whether this pocket represents the functional StheX-binding site, we next performed the structure determination of A167Y mutant enzyme. Crystal structure of A167Y mutant indeed indicated the decrease of the volume in the pocket due to the occupation of the planar and bulkier sidechain at this position (Figure 5B and 5C).

**Figure 5.**
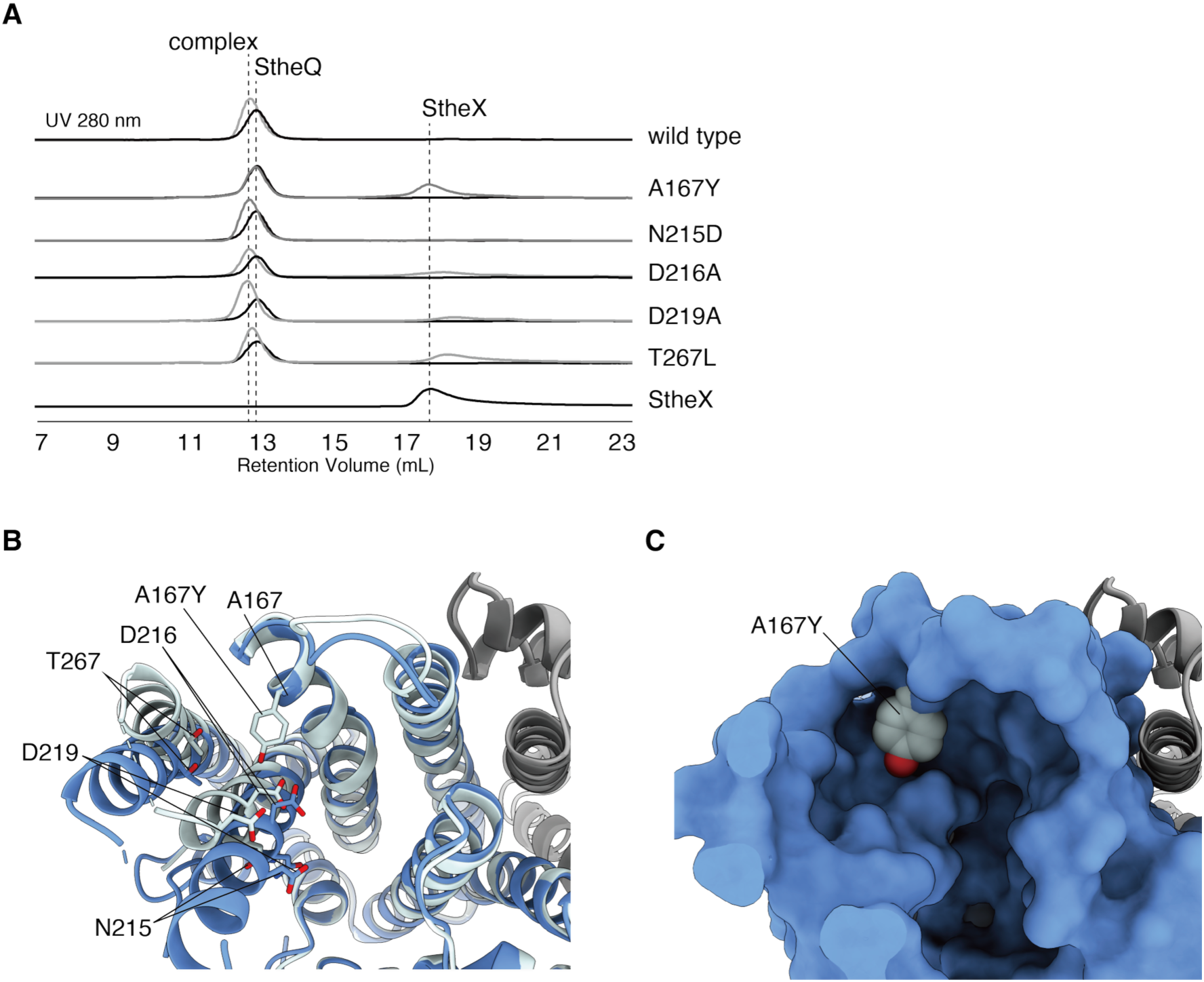
SEC elusion profiles of StheQ and StheQ mutants. (A) Chromatograms of SEC elution profiles of StheQ and StheQ mutants. The enzyme alone and the reaction mixture with StheX are shown as black and gray lines, respectively. (B and C) Crystal structure of A167Y mutant. (B) Superimposing of A167Y with StheQ wild type. The substituted residue is shown as a stick model. (C) Superimposing of sidechain of mutated Y167 onto the surface of StheQ wild type. The Tyr side chain is shown as a sphere model.

### Docking model of StheQ with StheX(60-69)

To further examine how the C-terminal region of StheX is accommodated in the putative peptide-binding pocket, we constructed the model structure. The structure-based analysis of peptide-protein interactions reported that most peptides do not induce conformational changes due to minimizing the entropic cost of binding.^41^ Thus, before docking simulation, we predicted the flexible region in StheX by ColabFold.^34^ The predicted structure of StheX was indicated that the region in the residues 1-59 was well-ordered with higher pLDDT values contrary whereas 60-69 residues exhibited <50 pLDDT values (Figure 6A). Thus, the residues 60-69, referred as StheX(60-69), may form the intrinsically disordered region. Furthermore, StheX(60-69) shares the similar sequence with the peptide in the matured ComX pheromone (Figure S1C). Then, we constructed a random coil structure of StheX(60-69) by CNS.^32^ For docking simulation, structure of StheQ-FSPP was slightly modified: α7 helix of molecule A was superimposed on the helix of molecule B, and then the residues 215-225 in the molecule A, which is disorder in the molecule B, replaced into the molecule B. Surprisingly, the symmetry mate of α8– α9 loop in StheQ also occupied the pocket comprising α6-α7 loop, and α6, α7, α8, and α9 helices (Figure S6), suggesting that the pocket was the candidate for accommodation of the ComX peptide. Thus, we performed docking simulation of StheX(60-69) against the surface lining A167 and T267 on the model structure of StheQ-FSPP by Autodock Vina 1.0.2.^33^ The docking study predicted that a part of StheX(60-69) was accommodated within the pocket (Figure 6B). In the docking model, N-terminus of StheX(60-69) was placed on the surface comprising α6-α7 loop, and α6, α7, α8, and α9 helices. C-terminus of StheX(60-69) was accommodated into the surface lining D82, E86, Q212, and N215. Main chain of G63 in StheX(60-69) was placed near the sidechain of A167 in StheQ. Main chains of I62, T64, and G66, and Oδ atom of T64 in StheX(60-69) interacted with β-phosphate. The substitution of Tyr for A167 occluded the pocket which accommodated the residues 62-64 of bound StheX(60-69) (Figure 6C–D). The substitution of Leu for T267 also partially occluded the binding site due to the replacement of the bulkier residue along with a loss of the hydrogen bond or salt bridge between T267 in StheQ and main chain of bound StheX(60-69). Therefore, this docking-based model supports the involvement of the predicted pocket in the accommodation of StheX. In Ord1, residues corresponding to Q212 and N215 of StheQ are positioned near the indole acceptor and have been proposed to contribute to aromatic prenylation through a proton-transfer relay, in which D219 initiates the reaction by promoting Q216-mediated abstraction of the N1 hydrogen from indole.^21^ By analogy, the docking model suggests that Q212 or N215 in StheQ may help position the side chain of W68 in StheX(60-69) at an appropriate distance from the C1 atom of the farnesyl group.

**Figure 6.**
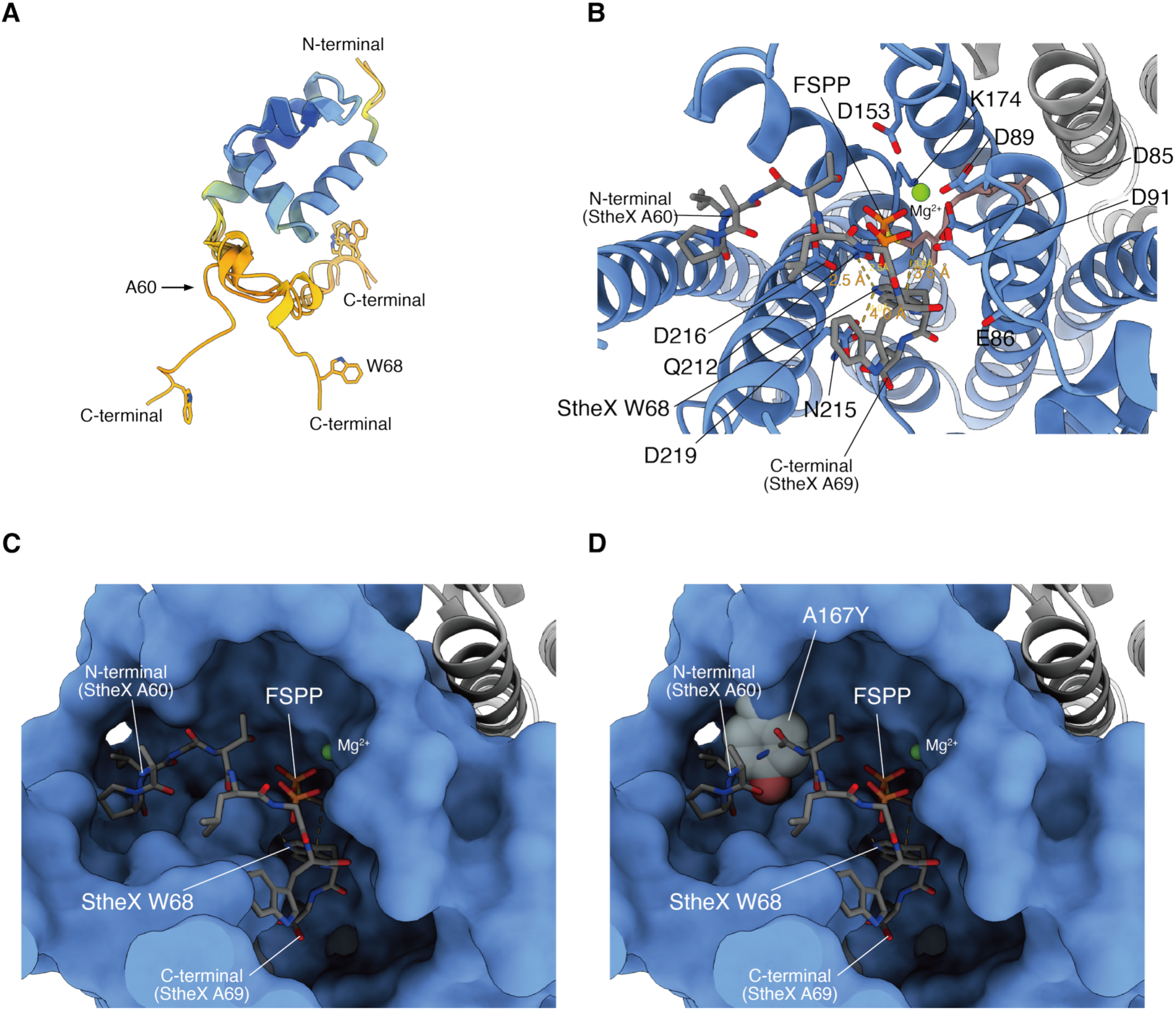
Proposed complex model of StheQ bound with StheX. (A) Three-dimensional model of StheX predicted using ColabFold. The values of pLDDT represented as a contintuous color scale. Blue, pLDDT value at 100; yellow, pLDDT value less than 50. Residues 1-59 forms the five predicted models are superimposed. (B) Docking model of StheQ bound with StheX(60-69). The StheX(60-69) peptide is shown as a gray stick model. The side chains of residues in FARM and pseudo-SARM and K174 and D153 coordinated with Mg^2+^ ion are also shown as stick models. Proposed key interaction between Oδ atom of N215 in StheQ and N1 atom of W68 in StheX and C1 atom in prenyl moiety and C3 atom in indole ring of W68 in StheX are shown as yellow dotted lines. (C and D) Binding mode of StheX(60-69) shown in (C) the surface model of StheQ and (D) the surface model of StheQ superimposed with A167Y mutant. The Tyr side chain is shown as a sphere model.

### Catalytic activities of the mutants

Structure analyses and the docking simulation identified several candidate residues that may contribute to the recognition and interaction of StheX. To investigate their functional roles, the prenylation activities of the mutants toward StheX(60-69) peptide were evaluated (Table 1). Surprisingly, although Q215 in Ord1, corresponding to Q212 in StheQ, was key residue for indole prenylation, Q212A mutant retained 94% activity relative to wild type. However, N215D mutant, which is located one helical pitch away from Q212 and positioned near the indole acceptor at 4.0 Å distance between Oδ atom of N215 and N1 atom of indole ring in the docking model, exhibited the 98% reduced prenylation activity. D216A and D219A mutants also exhibited markedly reduced activities with 96% and 83%, respectively. These results indicate that N215, together with D216 and D219, is important for efficient prenylation of StheX. Given their positions in the docking model, these residues may contribute to productive positioning and/or proton-transfer events involving the indole ring of W68 in StheX. Although A167Y markedly impaired complex formation with full-length StheX in the SEC assay, both A167Y and T267L mutants retained approximately 65% activity toward the C-terminal StheX(60-69) peptide. This difference may reflect distinct contributions of these residues to recognition of full-length StheX and to productive positioning of its C-terminal peptide. In the docking model, the C-terminal region of StheX containing W68 is positioned close to the catalytic center, whereas the A167 and T267 residues are predicted to interact with the N-terminal portion of the peptide. These results suggest that the mutations weaken binding of the peptide N-terminus thereby reducing the binding affinity and catalytic activity while maintaining an appropriate orientation of the peptide C-terminus (residues around W68) for catalysis. Thus, these docking and mutational analyses are compatible with the proposed involvement of this pocket in StheX recognition.

**Table 1.**
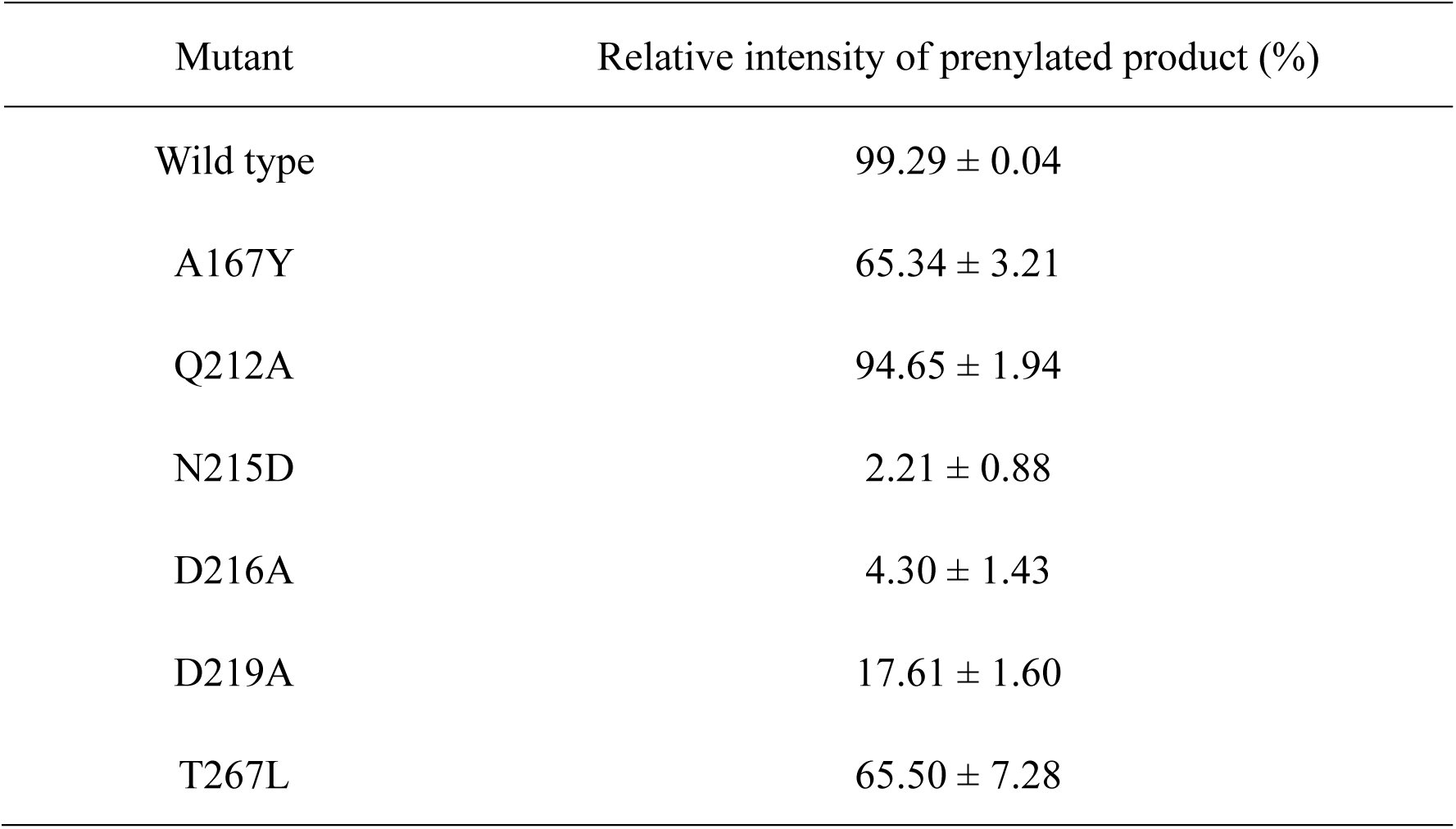
Summary of prenylated activities of mutant enzymes.

## DISCUSSION

The present study identifies StheQ as a peptide prenyltransferase acting on the cognate substrate StheX and expands the distribution of ComQ-like systems beyond the phylum *Firmicutes*. Although the biological role of the modified peptide remains unknown, the adjacent organization of *stheQ* and *stheX* genes, together with their functional relationship, suggests that StheQ and StheX may represent a previously unrecognized quorum-sensing-related peptide-modification system in *Chloroflexi*. Our structural and functional analyses suggest that the active-site architecture of StheQ more closely resembles that of the indole prenyltransferase Ord1 than that of canonical IPPS enzymes. Although StheQ retains the overall all-α-helical fold characteristic of the IPPS superfamily, the organization of its active site differs from that of chain-elongating IPPS enzymes and instead resembles that of aromatic prenyltransferases. This interpretation is further supported by comparison with Ord1.^21^ In particular, the architecture of the prenyl acceptor-binding pocket and the spatial arrangement of residues surrounding the pseudo-SARM region are consistent with adaptation for peptide-based indole prenylation rather than for small-molecule isoprenoid elongation.

Structural comparison with Ord1 highlights both similarities and distinctions. In both enzymes, the prenyl donor binds at the FARM region in coordination with a Mg^2+^ ion, and residues located one helical turn downstream of the pseudo-SARM are positioned near the predicted indole-binding site. In StheQ, FSPP occupies a pocket formed by helices α4, α6-α9, and the α6– α7 loop, in a manner analogous to that observed in Ord1. The coordination of a single Mg^2+^ ion at the FARM, together with interactions involving N215, is consistent with a catalytic model in which the prenyl donor is activated and stabilized in a geometry compatible with electrophilic aromatic substitution. Despite these similarities, StheQ differs from canonical IPPS enzymes in that no metal coordination is observed at the pseudo-SARM region, in contrast to the dual-metal coordination typically seen in chain-elongating IPPSs. This observation suggests that the pseudo-SARM in StheQ does not function as a second metal-binding motif in the present structures but may instead contribute to substrate positioning and catalytic specificity. The structural data therefore support the view that ComQ-family enzymes represent a distinct functional branch within the IPPS superfamily that has evolved to accommodate peptide substrates.

Mutational analyses further support the structural model of peptide accommodation. The A167Y substitution, which introduces a bulkier side chain into the predicted peptide-binding pocket, disrupts complex formation with StheX, consistent with steric occlusion of the binding cavity. The mutational analyses revealed that the residues in pseudo-SARM are important for prenylation activity toward StheX(60-69). Among these residues, N215 appears to play a particularly important role for the prenylation. Consistent with this interpretation, alterations near the pseudo-SARM region in ComQ have previously been shown to affect prenylation activity and substrate specificity.^40^ The SEC analysis, the A167Y structure, and mutational data support the presence of a peptide-binding pocket adjacent to the active site. Docking simulations further support a plausible structural model for how the flexible C-terminal region of StheX may be accommodated within this pocket and orient W68 toward the prenyl donor. N215 may also contribute to orienting the W68 side chain of StheX near the prenyl donor, although direct visualization of the enzyme–substrate complex will be required to confirm this arrangement.

Our structural observations are consistent with a plausible catalytic model in which StheQ coordinates the prenyl donor through the FARM-associated Mg²⁺ ion and positions the indole side chain of StheX for electrophilic substitution. The spatial arrangement of residues in the pseudo-SARM region suggests a role in substrate orientation and stabilization rather than canonical metal coordination observed in chain-elongating IPPS enzymes.

The active-site geometry is consistent with a catalytic architecture broadly analogous to that proposed for Ord1, in which activation of the prenyl donor precedes aromatic substitution. A schematic representation of this proposed catalytic model is shown in Figure 7. This model is intended to illustrate a plausible reaction pathway consistent with the structural data, rather than to define the precise sequence of proton-transfer events. Further biochemical and kinetic studies will be required to clarify whether the reaction proceeds through a discrete allylic carbocation intermediate and to determine the detailed proton-transfer mechanism.

**Figure 7.**
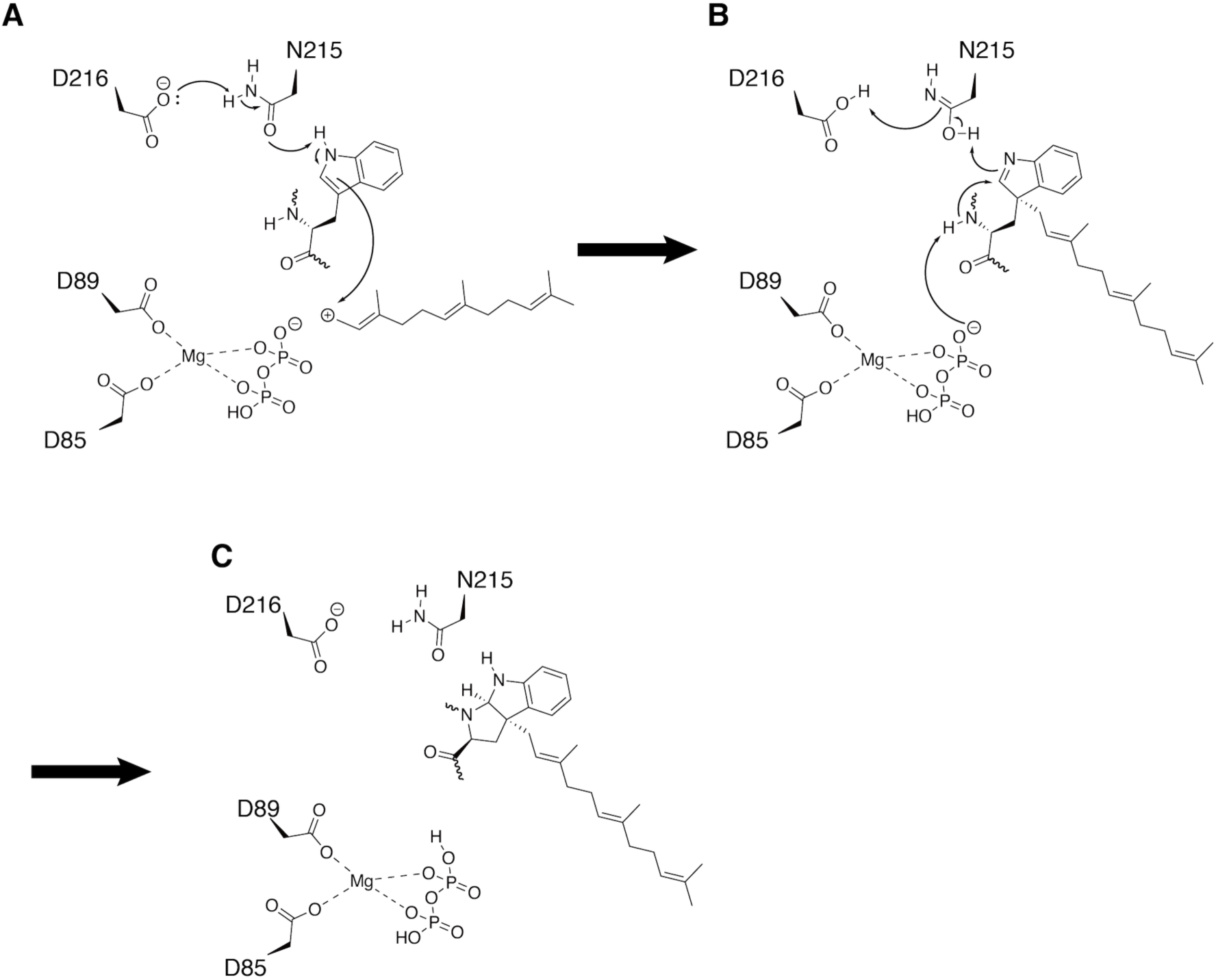
Proposed catalytic model for StheQ-mediated prenylation and cyclization. (A) A possible catalytic scenario consistent with the structural and mutational analyses. N215 may contribute to positioning of W68 and/or facilitate proton abstraction from N1 atom of its indole ring, followed by nucleophilic attack at the allylic carbocation generated from FPP. (B) In the proposed model, pyrophosphate may subsequently act as a base to promote deprotonation of the main-chain amide of W68, enabling intramolecular attack on the C2 atom of the indole ring. (C) Subsequent cyclization yields the Pro-like cyclic structure of the prenylated tryptophan residue, followed by product release.

These findings provide structural insight into the evolution of peptide-based indole prenylation within the IPPS superfamily and are consistent with the view that ComQ-family enzymes constitute a distinct functional branch specialized for peptide modification.

## Supporting information

Supporting Informations

## Author contributions

T.M and M.Okada designed the study; T.M., S.I., S.Y., A. K., R.T., A.S., and, M.Odagi prepared the recombinant proteins, and evaluated the enzymatic activities; T.M., and S.Y. performed structure determinations; T.M., S.I., and M.Okada, analyzed the data; T.M., S.I., and M.Okada wrote the paper, Y.K., H.M., and I.A supervised the project.

## ACKNOWLEDGMENT

The X-ray diffraction experiments were performed at the Photon Factory under proposals 14G504, 14G521, 14G530, and 15G007. The experiments at SPring-8 were performed under the proposal No. 2222 with support from the Platform Project for Supporting in Drug Discovery and Life Science Research (Platform for Drug Discovery, Informatics, and Structural Life Science) from the Japan Agency for Medical Research and Development (AMED).

