## Supporting Informations for "Identification and structural basis of a *Chloroflexus* protein with homology to *Bacillus* quorum sensing-related prenyltransferase"

### Supporting Information Figures

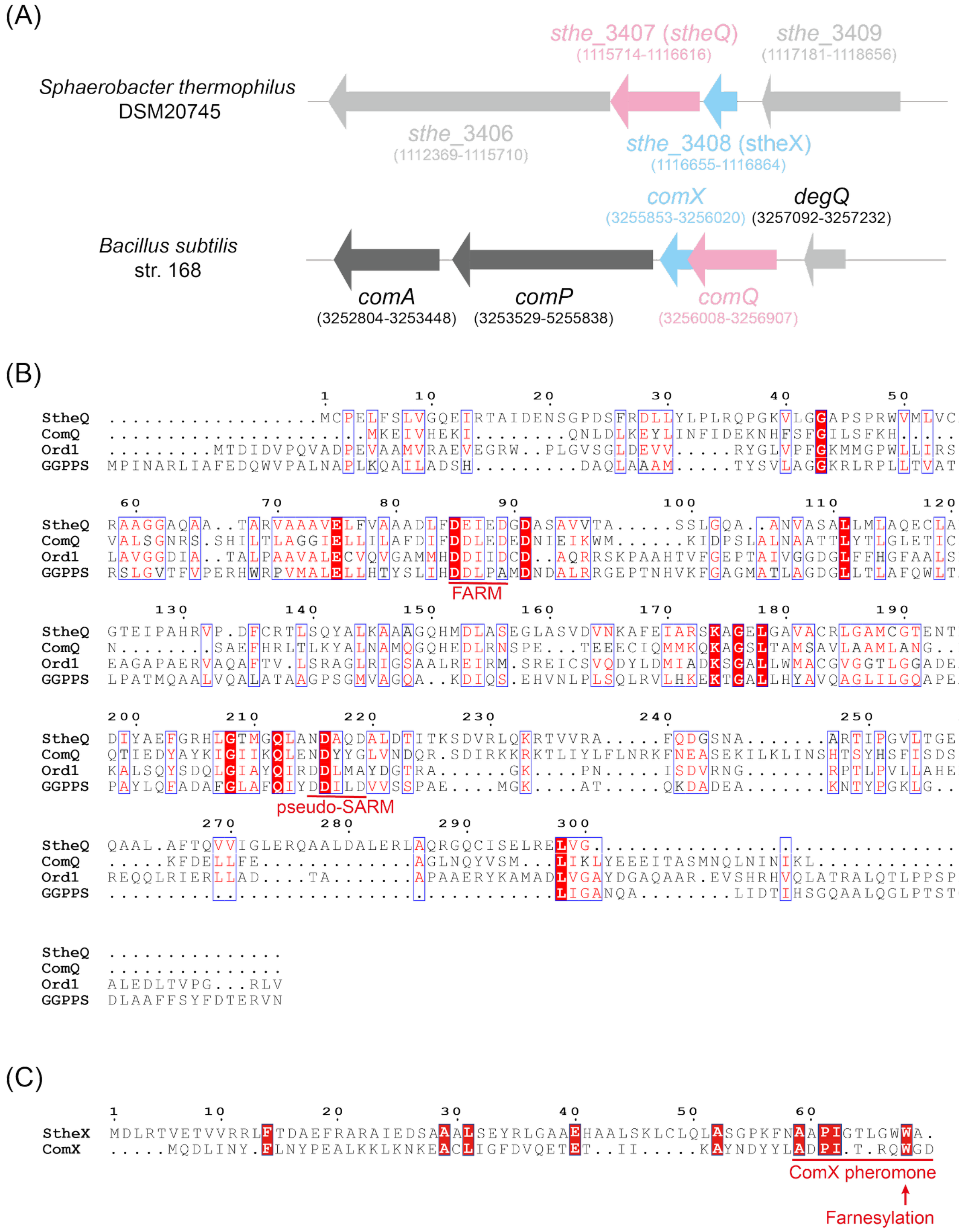


**Figure S1.** Gene loci of *stheXQ* and their amino acid sequences. **(A)** Comparison between the *comQXPA* gene cluster in the *B. subtilis* strain 168 and the corresponding locus including *stheQ* and *stheX* genes. (B) Sequence alignment of StheQ compared with ComQ (UniProt, P0DV09) from

*B. subtilis* strain RO-E-2, Ord1 (DDBJ, LC876682), and GGPPS (UniProt, Q03RR4). (C) Sequence alignment of StheX compared with ComX (UniProt, P45453) from *B. subtilis* strain 168.

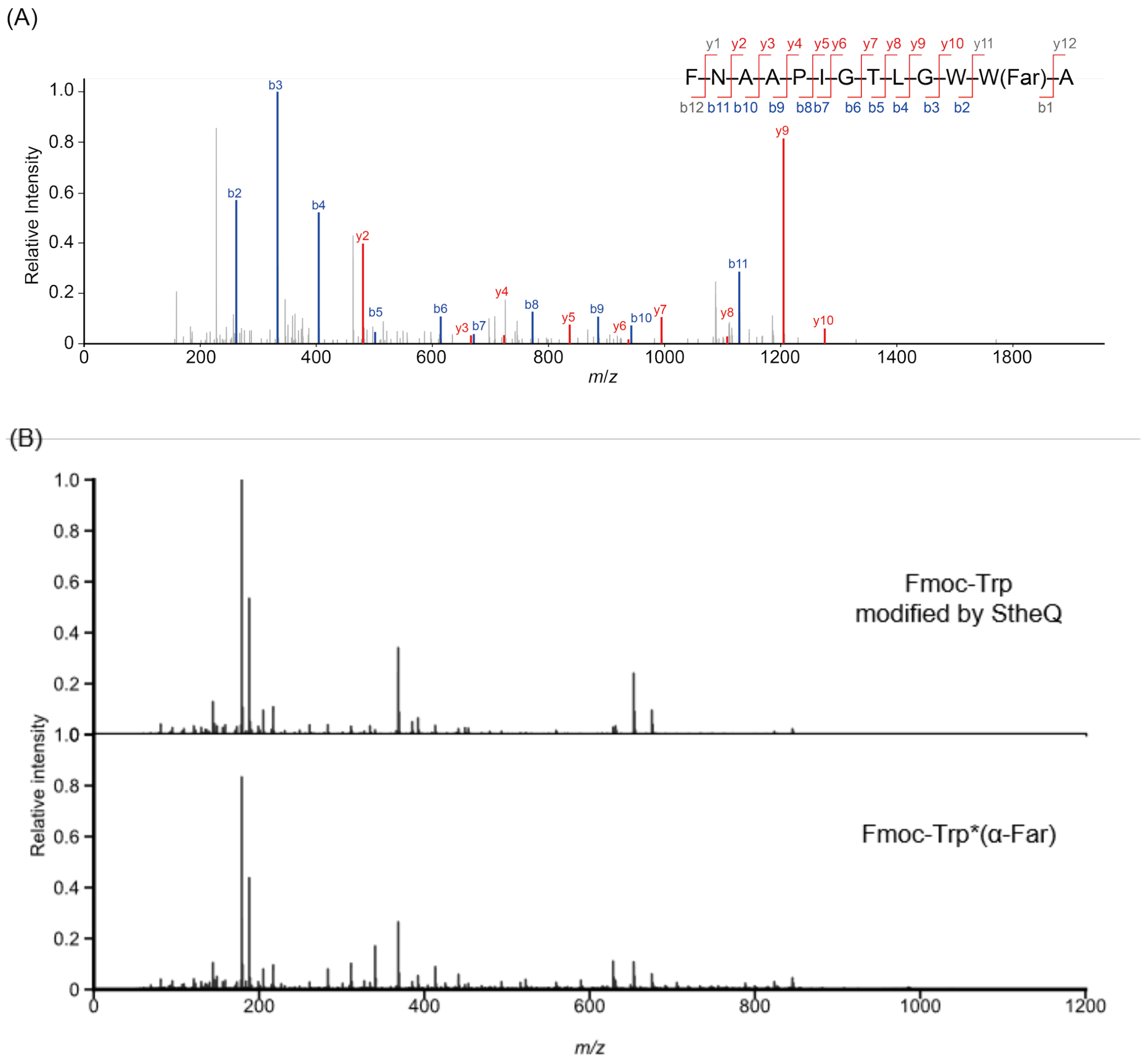


**Figure S2.** MS/MS spectra of farnesylated products. MS/MS spectra of (A) farnesylated StheX and (B) Fmoc-Trp. (A) MS/MS spectra of the tryptic peptide "FNAAPIGTLGWW(Far)A" (theoretical *m*/*z* = 804.45286, *z* = 2) ranging from 57–69 residues derived from StheX. The b- and y-ions are shown as blue and red lines, respectively. (B) MS/MS spectra of Fmoc-Trp treated with StheQ and the authentic standard of α-farnesylated Fmoc-Trp are shown.

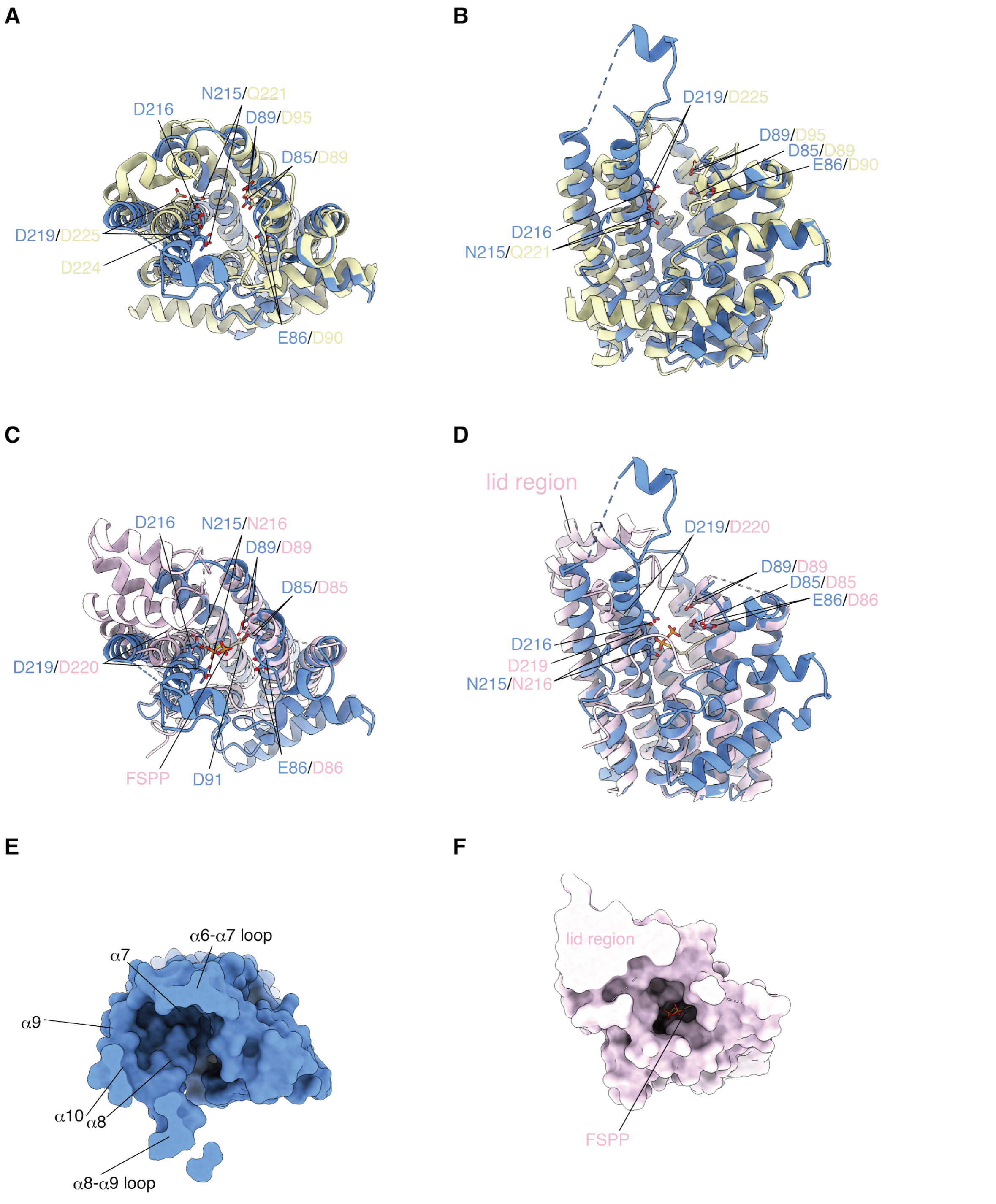


**Figure S3.** Structure comparisons of StheQ with other enzymes. (A–D) Superimposing of StheQ with (A and B) GGPPS (PDB 3PDE) and (C and D) Ord1 (PDB 9V4J). (A and C) Top view and (B and D) side view of structural comparison between protomers. The side chains of FARM,

SARM and pseudo-SARM are shown as stick model. Surface model of (E) StheQ and (F) Ord1 are shown. The lid region of Ord1 occupied at a part of wider pocket as seen in StheQ.

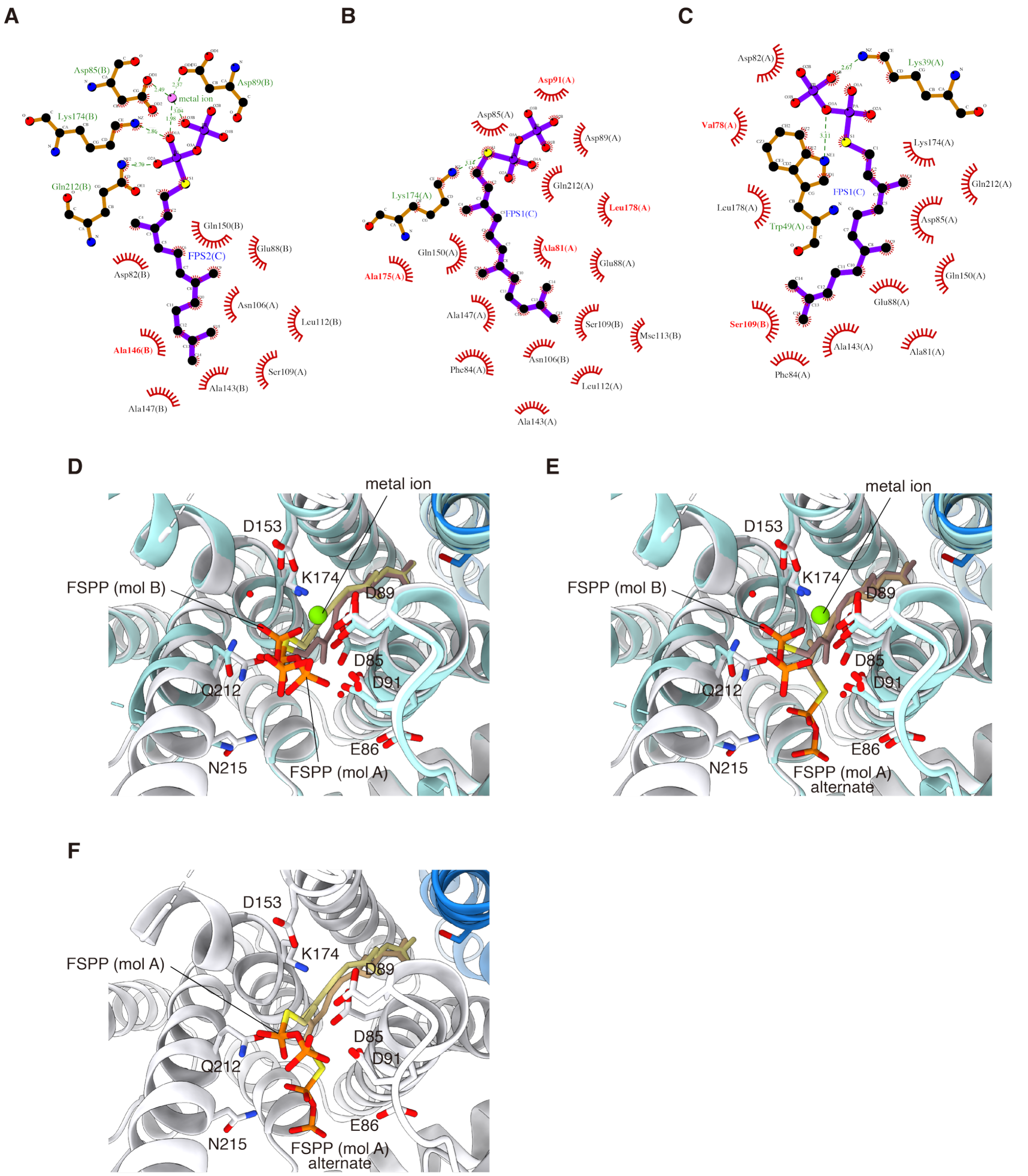


**Figure S4.** Binding modes of FSPP in the StheQ. (A–C) Illustrations of binding modes of FSPP in (A) molecule B, (B) molecule A, and (C) alternative mode shown in molecule A. These illustrations are drawn by LigPlot.[43] (D–E) Structural comparisons of binding modes between (D) molecule B and molecule A, (E) molecule B and alternative mode shown in molecule A, and (F) both binding modes shown in molecule A.

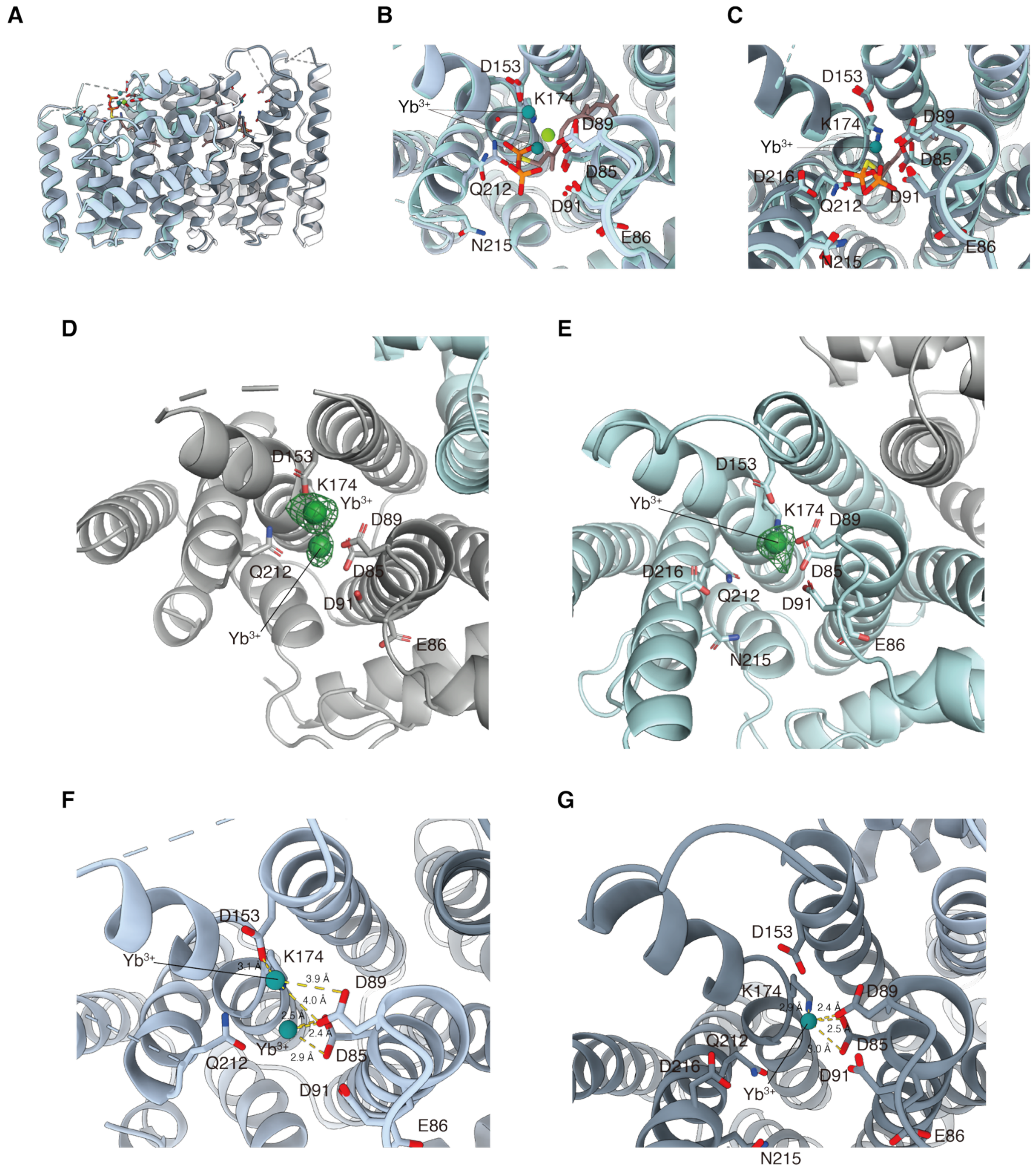


**Figure S5**. Coordination of metal ion. (A) Structural comparison between StheQ-FSPP and StheQ-$Yb^{3+}$ ion. (B and C) Closed view of FSPP binding site in (B) molecule B and molecule A. The complex structures of StheQ-FSPP and StheQ-$Yb^{3+}$ ion are shown as pale and dark cartoon models, respectively. The metal ions in the complex structures are shown as light and dark green spheres, respectively. (D and E) Crystal structures of molecule B and molecule A in StheQ-$Yb^{3+}$ ion. $F_o - F_c$ map at 5.0 σ is shown as a green mesh. (F and G) Coordination distances between $Yb^{3+}$ ion and sidechains of residues in (F) molecule B and (G) molecule A are shown as dashed lines.

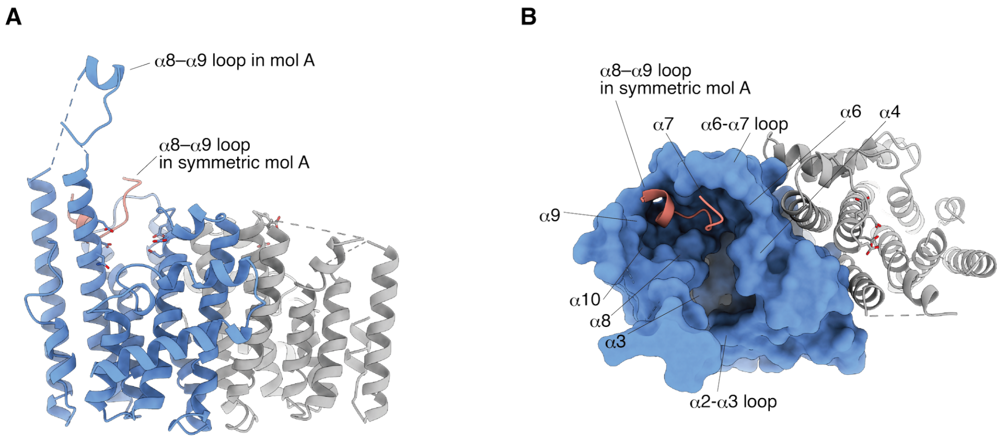


**Figure S6**. Interaction of α8–α9 loop in symmetry model into the pocket. Interaction between α8–α9 loop and molecule A in asymmetric unit is shown as (A) cartoon and (B) surface models, respectively. α8–α9 loop derived from symmetry model is shown as red cartoon.

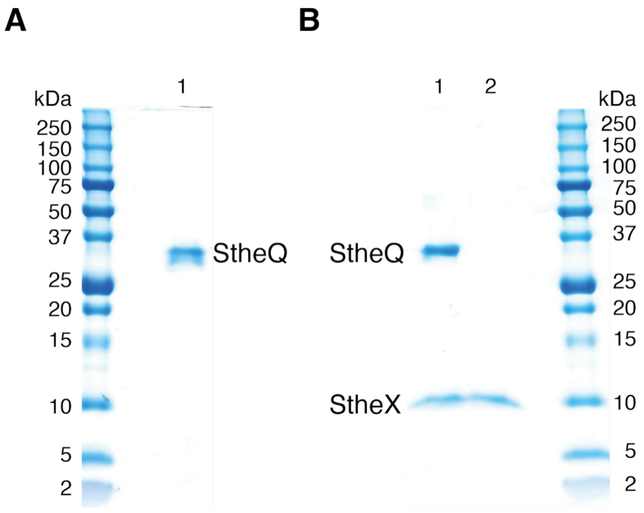


**Figure S7**. The SEC elution profiles of complex formation assays. (A and B) Tricine-SDS gels. (A) Lane 1, peak fraction from StheQ wild type in Figure 5A. (B) Lane 1, complex peak fraction from reaction mixture of StheQ and StheX in Figure 5A; lane 2, peak fraction from StheX in Figure 5A.

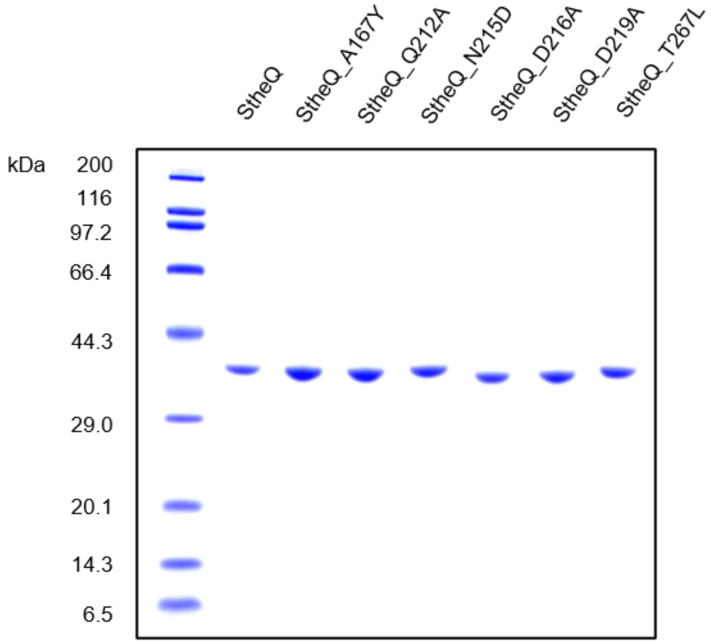


**Figure S8.** SDS-PAGE analysis of StheQ variants used for the activity assay (Table 1).

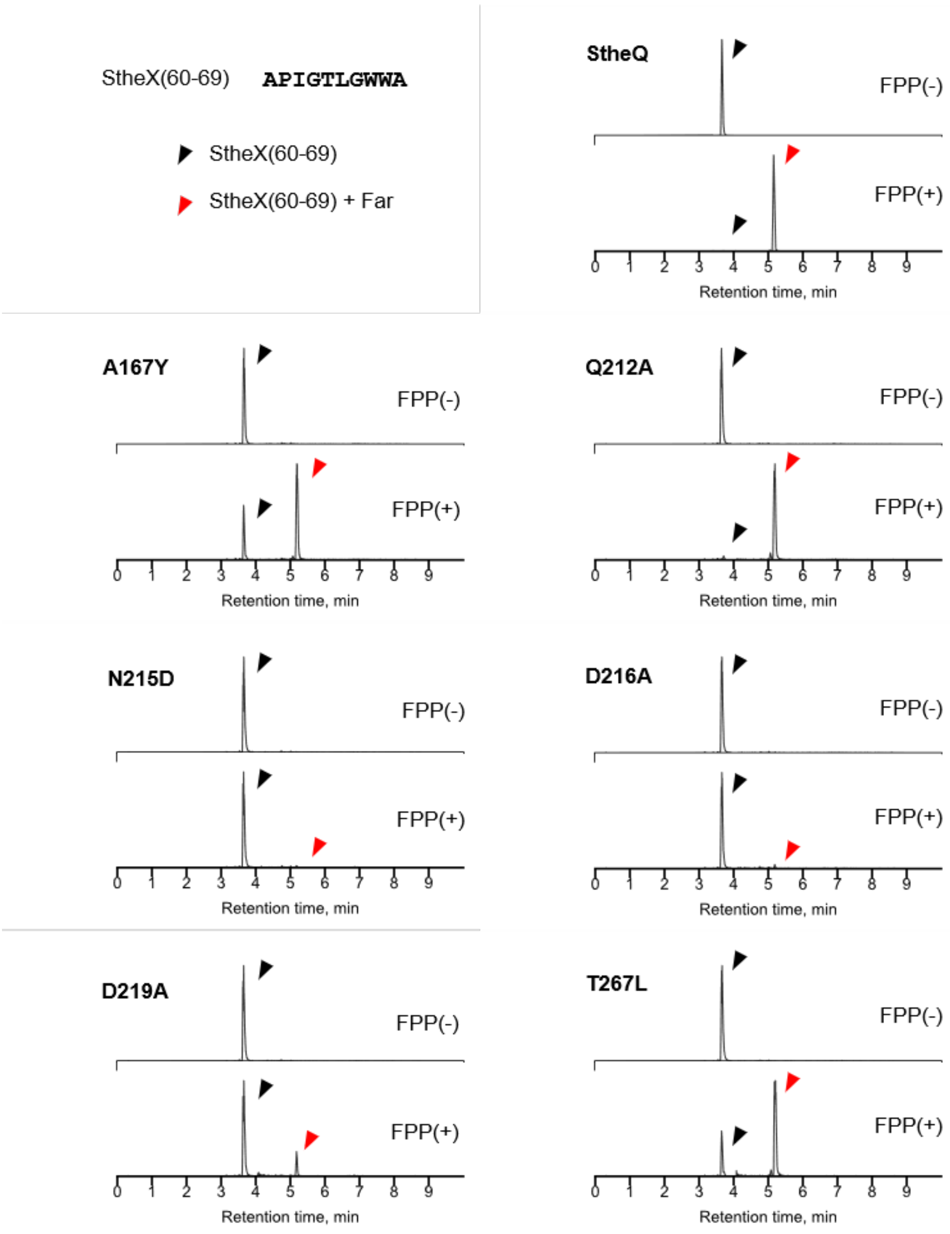


**Figure S9.** Farnesylation of StheX(60-69) mediated by StheQ mutants. Combined extracted ion chromatograms of substrates and their farnesylated products obtained from the enzyme variants treated with or without FPP are shown (Far: farnesyl). Unreacted substrates and farnesylated products are indicated by black and red arrows, respectively. Relative intensities of prenylated products are summarized in Table 1.

**Table S1.** List of the primer sequences used to generate StheQ mutants for activity assay.

| Mutant | Forward primer sequence | Reverse primer sequence |
|---|---|---|
| Q212A | ATGGGCGCGTTAGCCAACGATGCACAG | GGCTAACGCGCCCATCGTGCCCAGATG |
| N215D | TTAGCCGATGATGCACAGGATGCACTG | TGCATCATCGGCTAACTGGCCCATCGTG |
| D216A | GCCAACGCGGCACAGGATGCACTGGATAC | CTGTGCCGCGTTGGCTAACTGGCCCATC |
| D219A | GCACAGGCGGCACTGGATACCATCACC | CAGTGCCGCCTGTGCATCGTTGGCTAAC |
| A167Y | AACAAATATTTCGAAATTGCACGCTCC | TTCGAAATATTTGTTGACATCGACCGAAG |
| T267L | GCCTTTCTGCAGGTCGTGATTGGCCTG | GACCTGCAGAAAGGCTAAGGCAGCTTG |

**Table S2.** List of the primer sequences used to generate StheQ mutants for crystallization and SEC analyses.

| **Mutant** | **Forward primer sequence** | **Reverse primer sequence** |
|---|---|---|
| N215D | ACGATGGGGCAGTTGGCTGACGACGCACA | TGTGCGTCGTCAGCCAACTGCCCCATCGT |
| D216A | ATGGGGCAGTTGGCTAACGGCGCACAGGA | TCCTGTGCGCCGTTAGCCAACTGCCCCAT |
| D219A | TTGGCTAACGACGCACAGGGCGCTTTGGAT | ATCCAAAGCGCCCTGTGCGTCGTTAGCCAA |
| A167Y | TTTGCTCCTGGCAATCTCAAAGTACTTATTCACGTCGACCGAAGC | GCTTCGGTCGACGTGAATAAGTACTTTGAGATTGCCAGGAGCAAA |
| T267L | CCAATGACGACCTGGAGGAATGCGAGCGCGGC | CCAATGACGACCTGGAGGAATGCGAGCGCGGC |
